## Supplementry Materials for "Genetic Stability of *Mycobacterium smegmatis* under the Stress of First-Line Antitubercular Agents: Assessing Mutagenic Potential"

### Supplementary information

#### Abstract

The sustained success of *Mycobacterium tuberculosis* as a pathogen arises from its ability to persist within macrophages for extended periods and its limited responsiveness to antibiotics. Furthermore, the high incidence of resistance to the few available antituberculosis drugs is a significant concern, especially since the driving forces of the emergence of drug resistance are not clear. Drug-resistant strains of *Mycobacterium tuberculosis* can emerge through *de novo* mutations, however, mycobacterial mutation rates are low. To unravel the effects of antibiotic pressure on genome stability, we determined the genetic variability, phenotypic tolerance, DNA repair system activation, and dNTP pool upon treatment with current antibiotics using *Mycobacterium smegmatis*. Whole-genome sequencing revealed no significant increase in mutation rates after prolonged exposure to first-line antibiotics. However, the phenotypic fluctuation assay indicated rapid adaptation to antibiotics mediated by non-genetic factors. The upregulation of DNA repair genes, measured using qPCR, suggests that genomic integrity may be maintained through the activation of specific DNA repair pathways. Our results, indicating that antibiotic exposure does not result in *de novo* adaptive mutagenesis under laboratory conditions, do not lend support to the model suggesting antibiotic resistance development through drug pressure-induced microevolution.

#### Supplementary Figures

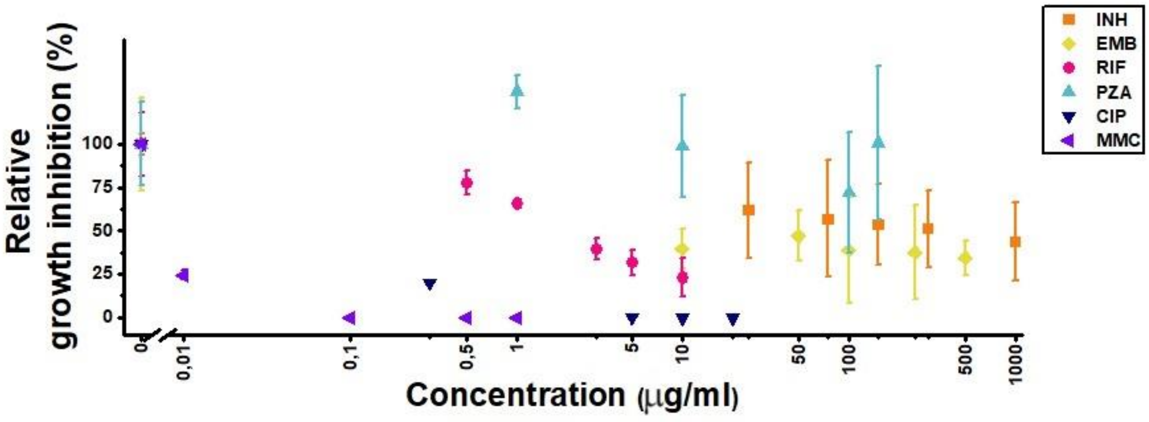


**Figure S1** Treatment optimization in liquid culture

Effects of treatment on *Mycobacterium smegmatis* mc$^2$-155 wild-type liquid cultures during the exponential growth phase. Relative growth inhibition is expressed as the % $OD_{600}$ ratio between treated and control liquid cultures for INH, EMB, RIF, and PZA (latter grown in pH 5.5 acidic media). For CIP and MMC, it is represented as the % ratio of CFUs following an 8-hour treatment.

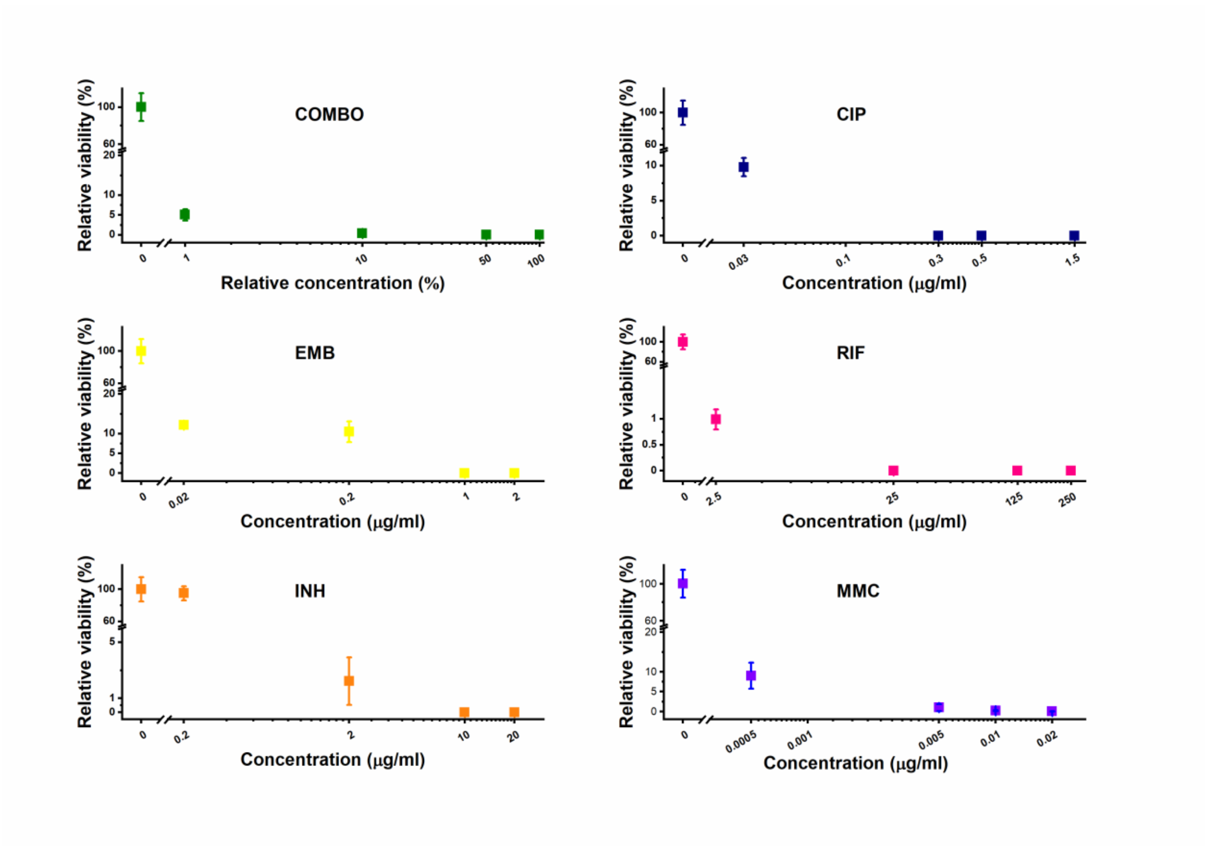


**Figure S2** Treatment optimization on agar plates

Relative viability is calculated based on the CFU counting of exponentially growing wild-type *M. smegmatis* cultures on control and selective plates containing the corresponding antibiotic at different concentrations. Incubation: 60 hours at 37°C.

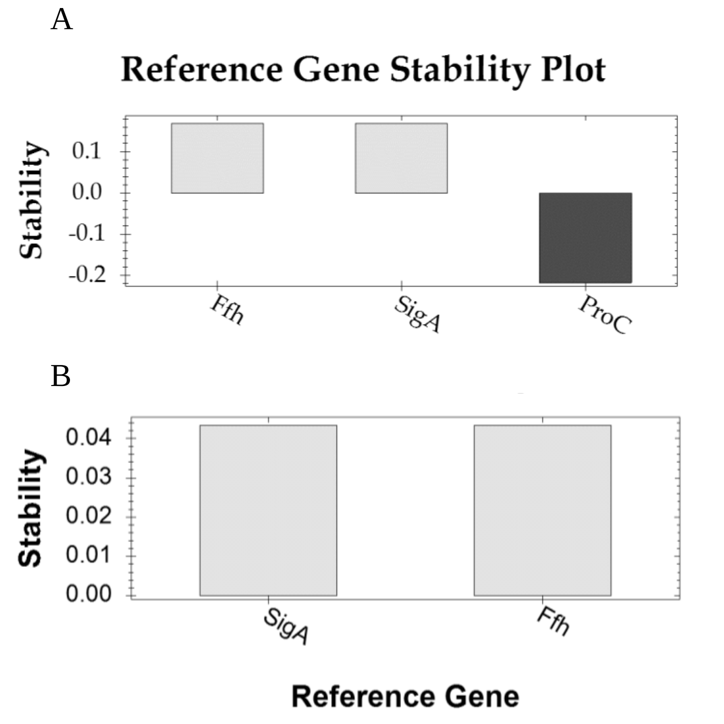


**Figure S3** Stability analysis of reference genes using the geNorm algorithm
A) Stability, denoted as 1/ln(M), where M represents the average pair-wise variation among the tested reference genes across all samples (including six treatments and controls, with three biological and three technical replicates each). M values were computed using the BioRad CFX Maestro$^{TM}$ software. Reference gene stability is considered acceptable within the 0-1 stability range and deemed unstable below 0, as determined by the software. ProC was excluded from the study due to its demonstrated instability. B) Pairwise comparison of the two accepted reference genes.

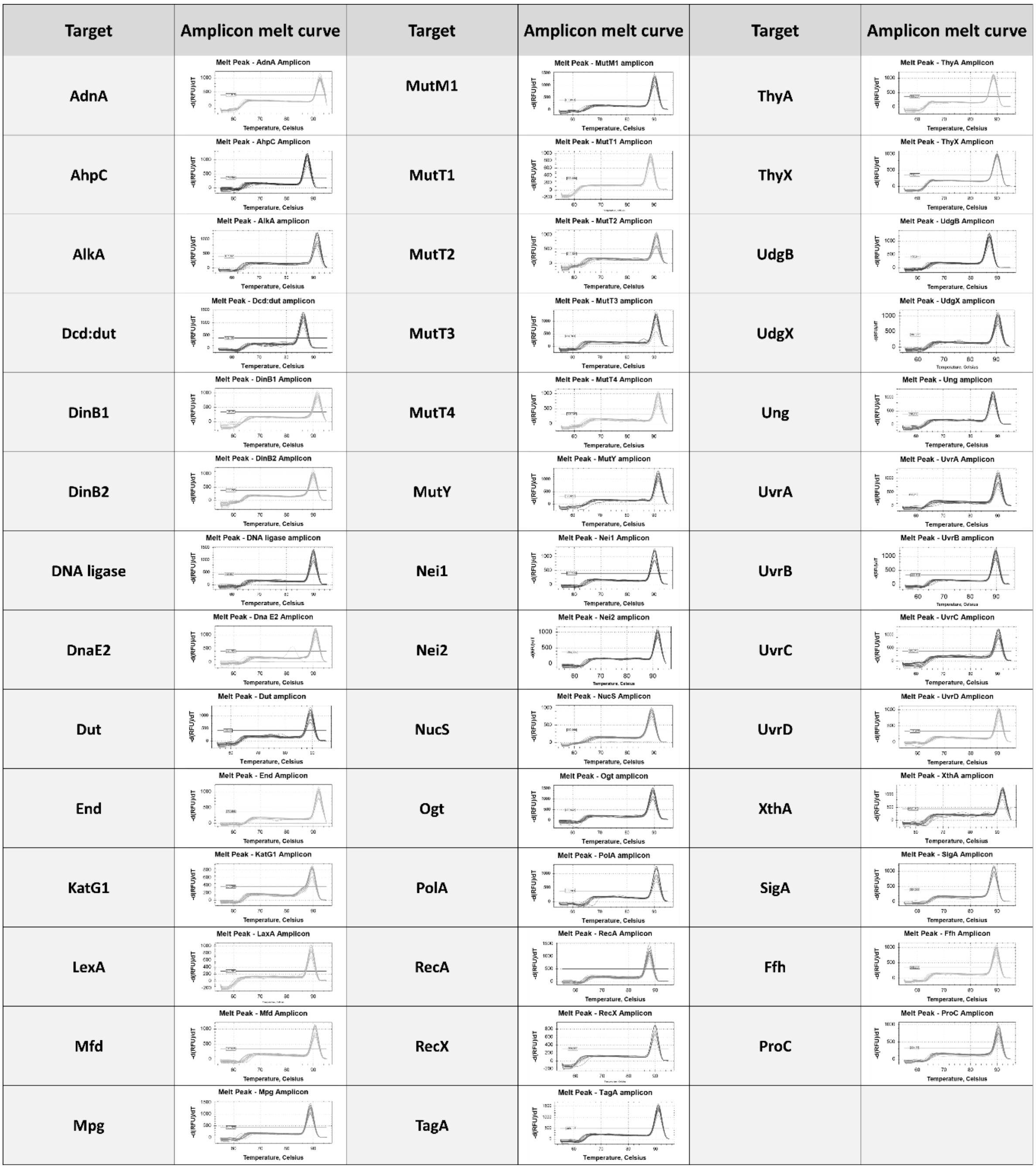


**Figure S4** Specificity assessment of the employed primers
Melt curve analysis of amplicons generated from target genes in the *Mycobacterium smegmatis* genome. The sequences of both forward and reverse primers are detailed in Table S2.

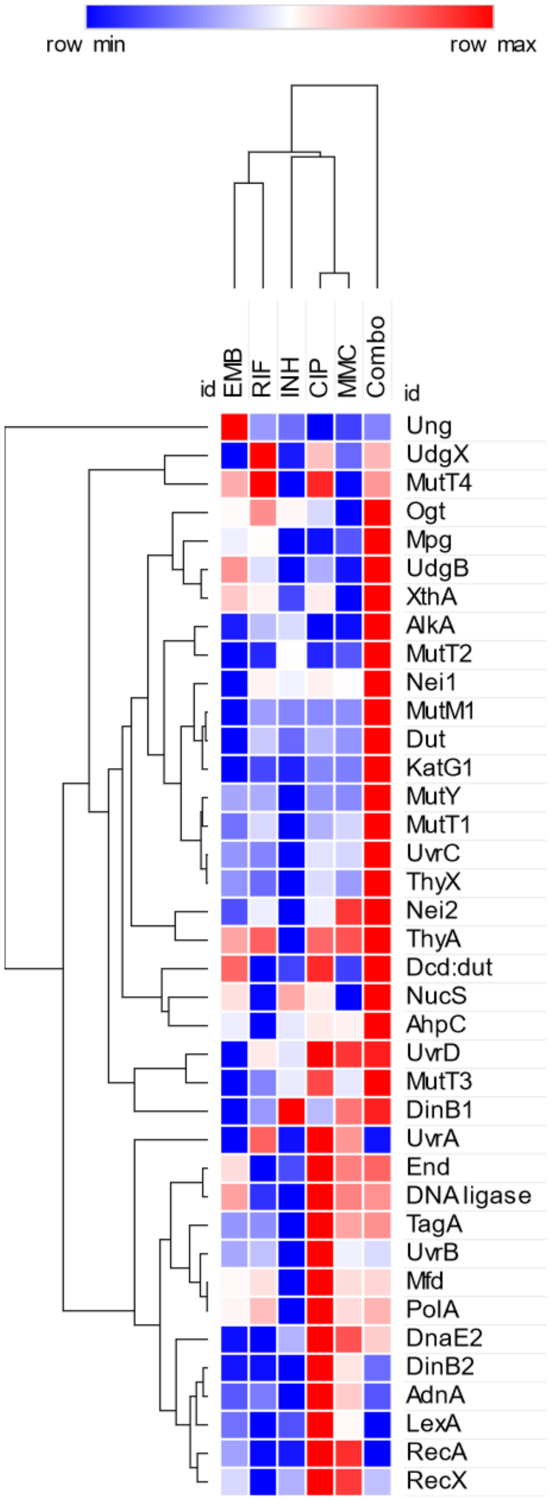


**Figure S5** Heatmap with clustering for gene expression changes upon treatment
The color scale is based on the mean relative gene expression changes compared to nontreated controls (fold change for upregulation and -1/fold change for downregulation). The color intensity in this heatmap is relative within each row and is not comparable to the color scale in Figure 3. Hierarchical clustering was performed using the one-minus Pearson correlation metric and the average linkage method. The figure was created using the Morpheus online heatmap generating tool (https://software.broadinstitute.org/morpheus/).

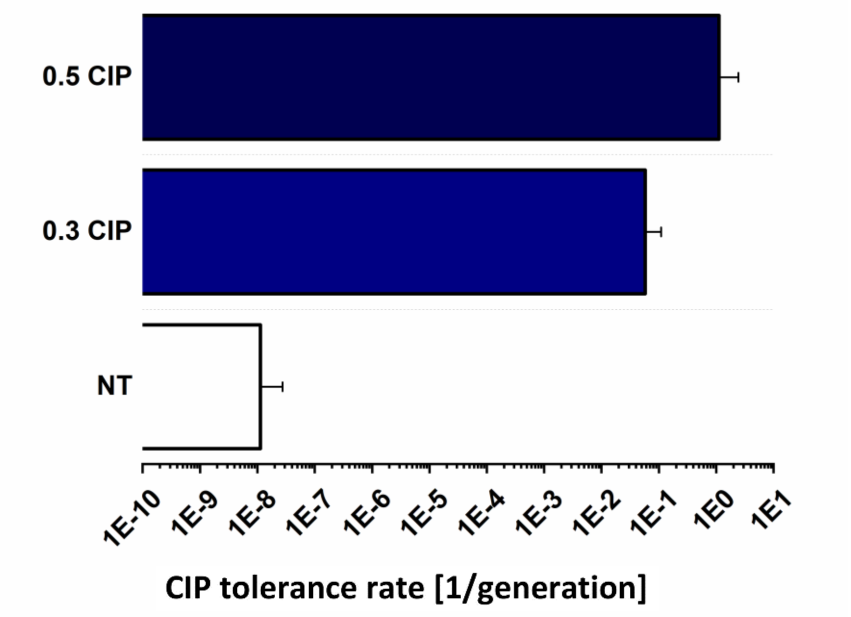


**Figure S6** CIP tolerance of *M. smegmatis* preincubated for 96h on CIP-containing plates sent for WGS
WT *M. smegmatis* colony was cultivated in LEMCO broth to reach OD=0.4-0.5, then striked onto plates containing 0.5 µg/ml CIP; 0.3 µg/ml CIP or onto non-selecting plates and grown for 4 days at 37 °C. Data bars represent the averages of three biological replicates each carried out in three technical replicates; error bars represent SE.

#### Supplementary Tables

**Table S1** Cell dimensions of *M. smegmatis* treated with different drugs

| | Control | COMBO | INH | EMB | RIF | MMC | CIP |
|---|---|---|---|---|---|---|---|
| Mean ($\mu m^3$) | 0.46 | 0.49 | 0.26 | 0.46 | 2.35 | 3.51 | 3.04 |
| SD | 0.25 | 0.18 | 0.13 | 0.21 | 0.88 | 1.71 | 1.14 |
| Mean length | 2.84 | 2.78 | 1.80 | 1.97 | 6.59 | 9.76 | 11.09 |
| SD | 0.92 | 0.70 | 0.54 | 0.78 | 2.41 | 4.58 | 4.04 |
| Mean width | 0.44 | 0.47 | 0.41 | 0.55 | 0.68 | 0.68 | 0.59 |
| SD | 0.08 | 0.05 | 0.07 | 0.14 | 0.09 | 0.11 | 0.10 |
| N | 212 | 100 | 84 | 91 | 95 | 118 | 137 |

**Table S2** Mutations detected from the lineages of each treatment

| Chromosome position | Sample | Reference | Mutation | AA mutation | Gene code | Uniprot protein name | Gene ontology (GO) | Experiment |
|---|---|---|---|---|---|---|---|---|
| 5214897 | cip_b | A | AG | Leu87 frameshift 148stop | MSMEG_5116 | Uncharacterized protein | N/A | Mutation accumulation (MA) |
| 3614832 | cip_b | C | T | Pro139Leu | MSMEG_3554 | N5,N10-methylene-tetrahydromethanopterin reductase | oxidoreductase activity, acting on paired donors, with incorporation or reduction of molecular oxygen [GO:0016705] | |
| 2208516 | cip_b | G | GA | Leu282 frameshift 283stop | MSMEG_2133 | Uncharacterized protein | N/A | |
| 5861538 | cip_c | G | GC | Leu168 frameshift | MSMEG_5792 | UPF0678 fatty acid-binding protein-like protein MSMEG_5792/MSMEI_5639 | intracellular transport [GO:0046907] | |
| 3415264 | cip_c | T | TC | Leu206 frameshift 257stop | MSMEG_3338 | Oxidoreductase, FAD/FMN-binding | FMN binding [GO:0010181]; oxidoreductase activity [GO:0016491] | |
| 2033295 | cip_c | A | AG | Leu72 frameshift 258stop | MSMEG_1954 | ABC1 family protein | N/A | |
| 1988098 | cip_c | A | AG | N/A | Intergenic region | intergenic | N/A | |
| 1533730 | inh_b | C | CTCG | Asp201_INSERTION | MSMEG_1431 | Cytochrome P450-terp (EC 1.14.-.-) | heme binding [GO:0020037]; iron ion binding [GO:0005506]; monooxygenase activity [GO:0004497]; oxidoreductase activity, acting on paired donors, with incorporation or reduction of molecular oxygen [GO:0016705] | |
| 994997 | inh_C | G | A | N/A | intergenic | | N/A | |
| 5777585 | inh_C | C | T | Val99Met | MSMEG_5688 | Regulatory protein, MarR | GO:0003700 DNA-binding transcription factor activity; GO:0006355 regulation of DNA-templated transcription | |
| 1508883 | mmc_a | C | G | Ala300Ala (neutral) | MSMEG_1407 | N/A | N/A | |
| 4598387 | mmc_a | C | G | Ala371Arg | MSMEG_4513 | Polyketide synthase | transferase activity, transferring acyl groups [GO:0016746] | |
| 6786854 | mmc_a | G | A | Trp104stop | MSMEG_6740 | 1-aminocyclopropane-1-carboxylate deaminase (EC 3.5.99.7) | 1-aminocyclopropane-1-carboxylate deaminase activity [GO:0008660]; pyridoxal phosphate binding [GO:0030170]; amine catabolic process [GO:0009310] | |
| 5313643 | mmc_c | C | T | N/A | intergenic | N/A | N/A | |
| 1865825 | mock_b | G | GC | Ala351 frameshift | MSMEG_1780 | Natural resistance-associated macrophage protein | metal ion transmembrane transporter activity; metal ion transport; membrane; | |
| 3722101 | mock_b | A | C | Asn185Thr | MSMEG_3656 | ABC transporter, permease/ATP-binding protein | | |
| 58213 | mock_c | T | TC | N/A | MSMEG_0037 | tRNA-Leu | N/A | |
| 4104684 | mock_c | T | C | Val70Ala | MSMEG_4033 | TetR-family protein transcriptional regulator | GO:0006350, Sequence-specific dna binding transcription factor activity, Regulation of transcription, dna-dependent | |
| 5118524 | mock_c | C | CG | Asp89 frameshift 143stop | MSMEG_5021 | Alcohol dehydrogenase, zinc-containing | Oxidoreductase activity, Zinc ion binding, Oxidation-reduction process | |
| 5217666 | mock_g | G | A | Thr200Thr (neutral) | MSMEG_5119 | L-glutamate gamma-semialdehyde dehydrogenase | Mitochondrial matrix, Oxidation-reduction process, Proline biosynthetic process, 1-pyrroline-5-carboxylate dehydrogenase activity | |
| 2970975 | mock_i | T | C | Arg155Gly | MSMEG_2908 | 2-Keto-3-deoxy-gluconate kinase | kinase activity [GO:0016301] | |
| 2970982 | mock_i | C | T | Arg153Glu | MSMEG_2908 | 2-Keto-3-deoxy-gluconate kinase | kinase activity [GO:0016301] | |
| 3306164 | mock_i | G | A | Glu1151Glu (neutral) | MSMEG_3225 | Ferredoxin-dependent glutamate synthase 1 (EC 1.4.7.1) | 3 iron, 4 sulfur cluster binding [GO:0051538]; glutamate synthase (ferredoxin) activity [GO:0016041]; metal ion binding [GO:0046872]; glutamate biosynthetic process [GO:0006537]; glutamine metabolic process [GO:0006541] | |
| 5805844 | mock_i | C | T | Val237Val (neutral) | MSMEG_5721 | Acetyl-CoA acetyltransferase | transferase activity, transferring acyl groups other than amino-acyl groups [GO:0016747] | |
| 4987517 | mock_j | G | A | Leu30Leu (neutral) | MSMEG_4890 | Alkyl hydroperoxide reductase AhpD (EC 1.11.1.28) (Alkylhydroperoxidase AhpD) | alkyl hydroperoxide reductase activity [GO:0008785]; hydroperoxide reductase activity [GO:0032843]; peroxidase activity [GO:0004601]; peroxiredoxin activity [GO:0051920]; response to oxidative stress [GO:0006979] | |
| 6406902 | mock_j | T | TG | N/A | | N/A | N/A | |
| 491016 | mock_k | C | T | N/A | intergenic | N/A | N/A | |
| 2287781 | mock_k | G | A | Gly199Asp | MSMEG_2207 | Beta-ketothiolase | transferase activity, transferring acyl groups other than amino-acyl groups [GO:0016747] | |
| 3438752 | rif_a | A | AC | Arg17 frameshift 175stop | MSMEG_3366 | Isonitrile hydratase, putative | N/A | |
| 5773058 | rif_a | C | T | Glu67Lys | MSMEG_5682 | Uncharacterized protein | integral component of membrane [GO:0016021] | |
| 6220187 | CIPB0.3 | G | T | Trp53Cys | MSMEG_6151 | Alpha/beta hydrolase fold-1 | epoxide hydrolase activity [GO:0004301] | Fluctuation assay with CIP treatment |

**Table S3** The nucleotide sequence and measured efficiency of primers used for the qPCR

| Target | Msm Gene ID | Sequence | Efficiency |
|---|---|---|---|
| AdnA_for | MSMEG_1941 | CGCAGTCCTACCGTTGC | 81.6 |
| AdnA_rev | | CTTCGGCGTGTGTGGAG | |
| AhpC_for | MSMEG_4753 | CCGACAAGCCCGAGAAG | 103 |
| AlhpC_rev | | GGAACGTGGACCGGATG | |
| AlkA_for | MSMEG_4925 | GCCTCCATCCGTCAGTTC | 83.5 |
| AlkaA_rev | | GCCGAACAATCCCTCGTAG | |
| Dcd:dut_for | MSMEG_0678 | CGGTTGGAGGGCAAGTC | 98.2 |
| Dcd:dut_rev | | CACAGCTGCCCGATCTTC | |
| DinB1_for | MSMEG_3172 | TCACGGTCAAGCTCAAGAAG | 83.2 |
| DinB1_rev | | AACCCAACGCCCACAAG | |
| DinB2_for | MSMEG_6443 | CCAGTTACGAGGCCAAGG | 81.2 |
| DinB2_rev | | GGGTGGTGTCGTGGAAG | |
| DNA ligase_for | MSMEG_2362 | GGAGGTCAAACGCAAGGG | 91.1 |
| DNA ligase_rev | | CAACTCGGTTCCGCACTC | |
| DnaE2_for | MSMEG_1633 | GCCTCGCTGGTGTTCTAC | 87.7 |
| DnaE2_rev | | AGGCTGGCATTCACGTC | |
| Dut_for | MSMEG_2765 | ATACCGCACGGAATGGTC | 97.1 |
| Dut_rev | | GTCTGCGGATCCAGGTTG | |
| End_for | MSMEG_1383 | AGCGCATCAAGTCCATCAC | 83 |
| End_rev | | GCCGTCTCGCAGATCAC | |
| Ffh_for | MSMEG_2430 | GAGCTCATCGGCATCCTC | 86.2 |
| Ffh_rev | | GGGCTGTGACCCTTGTC | |
| KatG1_for | MSMEG_6384 | GCCCATCGGAGAAGCTC | 96 |
| KatG1_rev | | CCTCGTCCAGCGGATTG | |
| LexA_for | MSMEG_2740 | GTTCCTGCTCAAGGTCGTC | 105 |
| LexA_rev | | CATGAGCCACACCTGACC | |
| Mfd_for | MSMEG_5423 | GAGCTCACCCGGTTCAC | 89 |
| Mfd_rev | | CTTGGTGGCGGTCTTCTC | |
| Mpg_for | MSMEG_3759 | GTGCGGAACTCGGTGATG | 89.6 |
| Mpg_rev | | GCTGTGCACTGTCGACTC | |
| MutM1_for | MSMEG_2419 | CAGGCGCGGAAAGTACC | 93 |
| MutM1_rev | | CGCTGATCGACGAAGCTC | |
| MutT1_for | MSMEG_2390 | GGTGGACAAGCTCGTATGG | 107 |
| MutT1_rev | | GCGATCGTCACCCTTGTAG | |
| MutT2_for | MSMEG_5148 | GGCTGTGGGAACTTCCTG | 86.7 |
| MutT2_rev | | TGTCATGGCGTCGTTGAG | |
| MutT3_for | MSMEG_0790 | CGTACACGACGGTGATCG | 92.1 |
| MutT3_rev | | GTAGGCGCTGCCAACTC | |
| MutT4_for | MSMEG_6927 | CAGGCGGTCTGGTCATC | 87.1 |
| MutT4_rev | | TGGATCCCGGTCTCCTC | |
| MutY_for | MSMEG_6083 | GGGAAAGCTCGGCTACC | 86.4 |
| MutY_rev | | GGAACGCTCGCCTGATAG | |
| Nei1_for | MSMEG_4683 | CATCGGCGCTCAGTACG | 99.7 |
| Nei1_rev | | CGGTCTGCGTGACTTGG | |

| | | | |
|---|---|---|---|
| Nei2_for | MSMEG_1756 | GCGGTACCGACATGGAC | 92.1 |
| Nei2_rev | | GAAACACAACTCGTTGCAGTAG | |
| NucS_for | MSMEG_4923 | CGCGCTACCTGGAACTG | 99.7 |
| NucS_rev | | GTACTCGTCGCTGTCCATTC | |
| Ogt_for | MSMEG_4928 | AGATCCCGTACGGACAGAC | 92.6 |
| Ogt_rev | | CCCATAACCCGTGAGACTTC | |
| PolA_for | MSMEG_3839 | GAGCTCACCCGGTTCAC | 83.9 |
| PolA_rev | | CTTGGTGGCGGTCTTCTC | |
| ProC_for | MSMEG_0943 | GCCCGGCGTACTTCTTC | 73.3 |
| ProC_rev | | CGGCGTTCACCTGATCC | |
| RecA_for | MSMEG_2723 | CGCGTCAAGGTCGTCAAG | 90.1 |
| RecA_rev | | ACCCTCGTAGGTGAACCAG | |
| RecX_for | MSMEG_2724 | CTCGAAACCCAGCTGACC | 89.9 |
| RecX_rev | | GAGCTCGACAGCCAAGG | |
| SigA_for | MSMEG_2758 | CATCTGCTGGAGGCGAAC | 87.7 |
| SigA_rev | | CTTGTAGCCCTTGGTGTAGTC | |
| TagA_for | MSMEG_5082 | GACTACCACGACACCGAATG | 91.4 |
| TagA_rev | | GGATCGAACCCGTGGAAC | |
| ThyA_for | MSMEG_2670 | GTCGGGTGAGCACATCG | 92.8 |
| ThyA_rev | | GGCGACGTAGAACTGGAAG | |
| ThyX_for | MSMEG_2683 | CGTACAGCTGATCGCCAAG | 94.2 |
| ThyX_rev | | CGTTGGTCGCGGTCTTC | |
| UdgB_for | MSMEG_5031 | ACGTTGACCACCGCATAC | 97.7 |
| UdgB_rev | | TTCGTTCCCGATCTGCTTG | |
| UdgX_for | MSMEG_0265 | CCCGGTGACAAAGAGGAC | 91.8 |
| UdgX_rev | | CTTGTGGATGCGTCGTTTG | |
| Ung_for | MSMEG_2399 | CCGTGGCAGATCAGGTG | 89.2 |
| Ung_rev | | TGTCGGGTAGGGATCCTG | |
| UvrA_for | MSMEG_3808 | CCTGGGCATCCGCAAAG | 94.6 |
| UvrA_rev | | CCCATCTCCTCACCGAGAC | |
| UvrB_for | MSMEG_3816 | GAGAAGGACAGCTCGATCAAC | 96.6 |
| UvrB_rev | | CTGACCCACCTGCAACTC | |
| UvrC_for | MSMEG_3078 | TCGATTTCTGCGACTTCCTG | 88.1 |
| UvrC_rev | | CCACGGCCTGTTTCTCC | |
| UvrD_for | MSMEG_5534 | GGGAGGACGGCATGTTC | 86.3 |
| UvrD_rev | | CGCGATTCCGGGTTGAG | |
| XthA_for | MSMEG_0829 | CCTACCACGGGCTCAAC | 80.7 |
| XthA_rev | | GCACGTAGAGGCTCCATAC | |

**Table S4** qPCR results

| Target | Treatment | Average fold change | +/- SEM | p |
|---|---|---|---|---|
| AdnA | CIP | 14.121 | 2.286 | 0.03887 |
| | NT | 1 | 0.199 | |
| AhpC | CIP | 0.752 | 0.07 | 0.59957 |
| | NT | 1 | 0.264 | |
| AlkA | CIP | 0.586 | 0.06 | 0.03689 |
| | NT | 1 | 0.507 | |
| Dcd:dut | CIP | 1.686 | 0.16 | 0.16879 |
| | NT | 1 | 0.222 | |
| DinB1 | CIP | 0.935 | 0.148 | 0.74175 |
| | NT | 1 | 0.478 | |
| DinB2 | CIP | 12.686 | 1.695 | 0.00768 |
| | NT | 1 | 0.2 | |
| DNA ligase | CIP | 3.171 | 0.156 | 0.05422 |
| | NT | 1 | 0.209 | |
| DnaE2 | CIP | 10.403 | 1.345 | 0.01572 |
| | NT | 1 | 0.2 | |
| Dut | CIP | 0.749 | 0.048 | 0.68244 |
| | NT | 1 | 0.347 | |
| End | CIP | 3.685 | 0.43 | 0.19237 |
| | NT | 1 | 0.213 | |
| KatG1 | CIP | 1.422 | 0.131 | 0.38096 |
| | NT | 1 | 0.228 | |
| LexA | CIP | 13.34 | 1.259 | 0.00578 |
| | NT | 1 | 0.2 | |
| Mfd | CIP | 3.833 | 0.267 | 0.03121 |
| | NT | 1 | 0.203 | |
| Mpg | CIP | 0.373 | 0.039 | 0.00387 |
| | NT | 1 | 0.411 | |
| MutM1 | CIP | 0.601 | 0.047 | 0.12714 |
| | NT | 1 | 0.274 | |
| MutT1 | CIP | 0.663 | 0.082 | 0.37964 |
| | NT | 1 | 0.342 | |
| MutT2 | CIP | 0.386 | 0.039 | 0.00986 |
| | NT | 1 | 0.705 | |
| MutT3 | CIP | 1.06 | 0.095 | 0.76275 |
| | NT | 1 | 0.442 | |
| MutT4 | CIP | 1.899 | 0.236 | 0.11622 |
| | NT | 1 | 0.252 | |
| MutY | CIP | 0.717 | 0.191 | 0.71034 |
| | NT | 1 | 0.35 | |
| Nei1 | CIP | 0.754 | 0.067 | 0.73054 |
| | NT | 1 | 0.249 | |

| | | | | |
|---|---|---|---|---|
| Nei2 | CIP | 0.769 | 0.066 | 0.47785 |
| | NT | 1 | 0.273 | |
| NucS | CIP | 1.028 | 0.106 | 0.90935 |
| | NT | 1 | 0.405 | |
| Ogt | CIP | 1.307 | 0.141 | 0.60338 |
| | NT | 1 | 0.264 | |
| PolA | CIP | 3.768 | 0.297 | 0.02211 |
| | NT | 1 | 0.204 | |
| RecA | CIP | 7.496 | 0.788 | 0.00229 |
| | NT | 1 | 0.198 | |
| RecX | CIP | 6.234 | 0.585 | 0.00557 |
| | NT | 1 | 0.204 | |
| TagA | CIP | 4.436 | 0.56 | 0.0077 |
| | NT | 1 | 0.203 | |
| ThyA | CIP | 0.396 | 0.032 | 0.0006 |
| | NT | 1 | 0.392 | |
| ThyX | CIP | 0.973 | 0.061 | 0.99138 |
| | NT | 1 | 0.231 | |
| UdgB | CIP | 1.036 | 0.119 | 0.85443 |
| | NT | 1 | 0.288 | |
| UdgX | CIP | 1.177 | 0.065 | 0.76245 |
| | NT | 1 | 0.324 | |
| Ung | CIP | 0.374 | 0.043 | 0.0091 |
| | NT | 1 | 0.846 | |
| UvrA | CIP | 1.982 | 0.159 | 0.07156 |
| | NT | 1 | 0.209 | |
| UvrB | CIP | 5.506 | 0.288 | 0.02087 |
| | NT | 1 | 0.202 | |
| UvrC | CIP | 0.874 | 0.071 | 0.20758 |
| | NT | 1 | 0.233 | |
| UvrD | CIP | 2.452 | 0.145 | 0.02696 |
| | NT | 1 | 0.202 | |
| XthA | CIP | 1.101 | 0.088 | 0.83909 |
| | NT | 1 | 0.229 | |
| AdnA | MMC | 7.902 | 1.464 | 0.05261 |
| | NT | 1 | 0.098 | |
| AhpC | MMC | 0.716 | 0.086 | 0.83256 |
| | NT | 1 | 0.115 | |
| AlkA | MMC | 0.677 | 0.105 | 0.69495 |
| | NT | 1 | 0.167 | |
| Dcd:dut | MMC | 0.933 | 0.144 | 0.9583 |
| | NT | 1 | 0.111 | |
| DinB1 | MMC | 1.179 | 0.335 | 0.78523 |
| | NT | 1 | 0.28 | |
| DinB2 | MMC | 7.513 | 1.366 | 0.0094 |
| | NT | 1 | 0.098 | |

| | | | | |
|---|---|---|---|---|
| DNA ligase | MMC | 1.991 | 0.357 | 0.34265 |
| | NT | 1 | 0.102 | |
| DnaE2 | MMC | 8.476 | 1.782 | 0.00215 |
| | NT | 1 | 0.099 | |
| Dut | MMC | 0.575 | 0.105 | 0.54121 |
| | NT | 1 | 0.133 | |
| End | MMC | 2.263 | 0.473 | 0.0107 |
| | NT | 1 | 0.119 | |
| KatG1 | MMC | 1.018 | 0.124 | 0.86562 |
| | NT | 1 | 0.117 | |
| LexA | MMC | 6.272 | 1.515 | 0.19399 |
| | NT | 1 | 0.099 | |
| Mfd | MMC | 1.417 | 0.229 | 0.4306 |
| | NT | 1 | 0.104 | |
| Mpg | MMC | 0.687 | 0.097 | 0.11023 |
| | NT | 1 | 0.312 | |
| MutM1 | MMC | 0.633 | 0.078 | 0.64028 |
| | NT | 1 | 0.123 | |
| MutT1 | MMC | 0.868 | 0.127 | 0.90405 |
| | NT | 1 | 0.126 | |
| MutT2 | MMC | 0.602 | 0.07 | 0.5445 |
| | NT | 1 | 0.162 | |
| MutT3 | MMC | 0.725 | 0.118 | 0.8164 |
| | NT | 1 | 0.23 | |
| MutT4 | MMC | 0.807 | 0.127 | 0.84788 |
| | NT | 1 | 0.154 | |
| MutY | MMC | 0.659 | 0.192 | 0.68625 |
| | NT | 1 | 0.248 | |
| Nei1 | MMC | 0.659 | 0.115 | 0.7256 |
| | NT | 1 | 0.175 | |
| Nei2 | MMC | 1.373 | 0.183 | 0.79159 |
| | NT | 1 | 0.122 | |
| NucS | MMC | 0.902 | 0.208 | 0.98398 |
| | NT | 1 | 0.194 | |
| Ogt | MMC | 0.72 | 0.1 | 0.9951 |
| | NT | 1 | 0.182 | |
| PolA | MMC | 1.353 | 0.233 | 0.2856 |
| | NT | 1 | 0.108 | |
| RecA | MMC | 6.686 | 0.934 | 0.00115 |
| | NT | 1 | 0.098 | |
| RecX | MMC | 5.343 | 0.81 | 0.09002 |
| | NT | 1 | 0.099 | |
| TagA | MMC | 1.931 | 0.517 | 0.29158 |
| | NT | 1 | 0.108 | |
| ThyA | MMC | 0.614 | 0.089 | 0.65887 |
| | NT | 1 | 0.145 | |

| | | | | |
|---|---|---|---|---|
| ThyX | MMC | 0.56 | 0.067 | 0.66447 |
| | NT | 1 | 0.143 | |
| UdgB | MMC | 0.989 | 0.203 | 0.89605 |
| | NT | 1 | 0.178 | |
| UdgX | MMC | 0.854 | 0.192 | 0.94767 |
| | NT | 1 | 0.122 | |
| Ung | MMC | 0.505 | 0.116 | 0.42393 |
| | NT | 1 | 0.369 | |
| UvrA | MMC | 1.057 | 0.203 | 0.82833 |
| | NT | 1 | 0.157 | |
| UvrB | MMC | 2.065 | 0.278 | 0.21923 |
| | NT | 1 | 0.136 | |
| UvrC | MMC | 0.769 | 0.091 | 0.79579 |
| | NT | 1 | 0.133 | |
| UvrD | MMC | 1.604 | 0.218 | 0.16877 |
| | NT | 1 | 0.102 | |
| XthA | MMC | 0.572 | 0.156 | 0.37319 |
| | NT | 1 | 0.21 | |
| AdnA | COMBO | 1.151 | 0.398 | 0.99589 |
| | NT | 1 | 1.787 | |
| AhpC | COMBO | 1.263 | 0.253 | 0.40096 |
| | NT | 1 | 1.069 | |
| AlkA | COMBO | 8.576 | 1.051 | 0.03138 |
| | NT | 1 | 0.155 | |
| Dcd:dut | COMBO | 1.972 | 0.305 | 0.21211 |
| | NT | 1 | 0.549 | |
| DinB1 | COMBO | 2.095 | 0.588 | 0.69326 |
| | NT | 1 | 1.429 | |
| DinB2 | COMBO | 3.525 | 0.738 | 0.02779 |
| | NT | 1 | 0.858 | |
| DNA ligase | COMBO | 1.85 | 0.271 | 0.33174 |
| | NT | 1 | 0.789 | |
| DnaE2 | COMBO | 5.629 | 1.384 | 0.00555 |
| | NT | 1 | 0.303 | |
| Dut | COMBO | 2.309 | 0.43 | 0.21693 |
| | NT | 1 | 0.274 | |
| End | COMBO | 2.559 | 0.399 | 0.05372 |
| | NT | 1 | 0.502 | |
| KatG1 | COMBO | 17.166 | 2.578 | 0.01456 |
| | NT | 1 | 0.115 | |
| LexA | COMBO | 0.851 | 0.167 | 0.66476 |
| | NT | 1 | 1.692 | |
| Mfd | COMBO | 1.505 | 0.344 | 0.25422 |
| | NT | 1 | 0.647 | |
| Mpg | COMBO | 5.55 | 1.388 | 0.00063 |
| | NT | 1 | 0.184 | |

| | | | | |
|---|---|---|---|---|
| MutM1 | COMBO | 4.033 | 0.577 | 0.04938 |
| | NT | 1 | 0.205 | |
| MutT1 | COMBO | 1.859 | 0.373 | 0.11133 |
| | NT | 1 | 0.599 | |
| MutT2 | COMBO | 6.075 | 0.676 | 0.02665 |
| | NT | 1 | 0.263 | |
| MutT3 | COMBO | 1.932 | 0.314 | 0.09768 |
| | NT | 1 | 1.018 | |
| MutT4 | COMBO | 1.129 | 0.304 | 0.80645 |
| | NT | 1 | 1.525 | |
| MutY | COMBO | 2.661 | 0.906 | 0.20534 |
| | NT | 1 | 0.429 | |
| Nei1 | COMBO | 2.621 | 0.541 | 0.117 |
| | NT | 1 | 0.545 | |
| Nei2 | COMBO | 2.029 | 0.413 | 0.3601 |
| | NT | 1 | 0.6 | |
| NucS | COMBO | 2.894 | 0.467 | 0.10342 |
| | NT | 1 | 0.335 | |
| Ogt | MMC | 4.981 | 0.553 | 0.0046 |
| | NT | 1 | 0.182 | |
| PolA | MMC | 1.793 | 0.285 | 0.19061 |
| | NT | 1 | 1.276 | |
| RecA | COMBO | 0.707 | 0.062 | 0.65932 |
| | NT | 1 | 0.871 | |
| RecX | COMBO | 1.323 | 0.322 | 0.50598 |
| | NT | 1 | 0.624 | |
| TagA | COMBO | 2.222 | 0.341 | 0.06179 |
| | NT | 1 | 1.072 | |
| ThyA | COMBO | 1.671 | 0.274 | 0.20173 |
| | NT | 1 | 1.409 | |
| ThyX | COMBO | 2.409 | 0.356 | 0.15855 |
| | NT | 1 | 1.014 | |
| UdgB | COMBO | 5.429 | 0.486 | 0.00956 |
| | NT | 1 | 0.285 | |
| UdgX | COMBO | 1.292 | 0.21 | 0.61771 |
| | NT | 1 | 1.21 | |
| Ung | COMBO | 0.809 | 0.269 | 0.78984 |
| | NT | 1 | 2.765 | |
| UvrA | COMBO | 0.95 | 0.115 | 0.70907 |
| | NT | 1 | 1.916 | |
| UvrB | COMBO | 1.789 | 0.28 | 0.13036 |
| | NT | 1 | 0.78 | |
| UvrC | COMBO | 2.551 | 0.259 | 0.1212 |
| | NT | 1 | 0.264 | |
| UvrD | COMBO | 1.987 | 0.251 | 0.06771 |
| | NT | 1 | 0.38 | |

| | | | | |
|---|---|---|---|---|
| XthA | COMBO | 3.548 | 0.923 | 0.00032 |
| | NT | 1 | 0.541 | |
| AdnA | INH | 0.507 | 0.307 | 0.7943 |
| | NT | 1 | 0.773 | |
| AhpC | INH | 0.416 | 0.159 | 0.19812 |
| | NT | 1 | 0.298 | |
| AlkA | INH | 2.024 | 0.778 | 0.40531 |
| | NT | 1 | 0.142 | |
| Dcd:dut | INH | 0.711 | 0.363 | 0.33358 |
| | NT | 1 | 0.198 | |
| DinB1 | INH | 1.855 | 1.13 | 0.72874 |
| | NT | 1 | 0.208 | |
| DinB2 | INH | 0.833 | 0.45 | 0.5528 |
| | NT | 1 | 0.291 | |
| DNA ligase | INH | 0.255 | 0.118 | 0.1147 |
| | NT | 1 | 0.607 | |
| DnaE2 | INH | 2.006 | 0.771 | 0.87827 |
| | NT | 1 | 0.172 | |
| Dut | INH | 0.341 | 0.141 | 0.07244 |
| | NT | 1 | 0.496 | |
| End | INH | 0.618 | 0.262 | 0.3347 |
| | NT | 1 | 0.272 | |
| KatG1 | INH | 0.252 | 0.106 | 0.02727 |
| | NT | 1 | 0.691 | |
| LexA | INH | 0.846 | 0.345 | 0.87991 |
| | NT | 1 | 0.225 | |
| Mfd | INH | 0.434 | 0.164 | 0.17385 |
| | NT | 1 | 0.342 | |
| Mpg | INH | 0.344 | 0.257 | 0.85723 |
| | NT | 1 | 0.763 | |
| MutM1 | INH | 0.444 | 0.179 | 0.19173 |
| | NT | 1 | 0.376 | |
| MutT1 | INH | 0.23 | 0.092 | 0.12512 |
| | NT | 1 | 0.659 | |
| MutT2 | INH | 1.074 | 0.446 | 0.45787 |
| | NT | 1 | 0.232 | |
| MutT3 | INH | 0.565 | 0.233 | 0.27115 |
| | NT | 1 | 0.407 | |
| MutT4 | INH | 0.6 | 0.266 | 0.52969 |
| | NT | 1 | 0.278 | |
| MutY | INH | 0.249 | 0.108 | 0.13517 |
| | NT | 1 | 0.668 | |
| Nei1 | INH | 0.428 | 0.176 | 0.10021 |
| | NT | 1 | 0.541 | |
| Nei2 | INH | 0.178 | 0.087 | 0.00101 |
| | NT | 1 | 0.731 | |

| | | | | |
|---|---|---|---|---|
| NucS | INH | 1.184 | 0.574 | 0.8894 |
| | NT | 1 | 0.228 | |
| Ogt | INH | 1.436 | 0.672 | 0.51484 |
| | NT | 1 | 0.142 | |
| PolA | INH | 0.403 | 0.164 | 0.18206 |
| | NT | 1 | 0.339 | |
| RecA | INH | 0.726 | 0.278 | 0.53931 |
| | NT | 1 | 0.197 | |
| RecX | INH | 0.797 | 0.305 | 0.86511 |
| | NT | 1 | 0.195 | |
| TagA | INH | 0.221 | 0.094 | 0.08878 |
| | NT | 1 | 0.55 | |
| ThyA | INH | 0.04 | 0.021 | 0.01628 |
| | NT | 1 | 6.6 | |
| ThyX | INH | 0.208 | 0.106 | 0.18768 |
| | NT | 1 | 0.803 | |
| UdgB | INH | 0.622 | 0.24 | 0.19842 |
| | NT | 1 | 0.146 | |
| UdgX | INH | 0.371 | 0.143 | 0.35299 |
| | NT | 1 | 0.262 | |
| Ung | INH | 0.516 | 0.196 | 0.33083 |
| | NT | 1 | 0.236 | |
| UvrA | INH | 0.729 | 0.301 | 0.89194 |
| | NT | 1 | 0.212 | |
| UvrB | INH | 0.749 | 0.299 | 0.99766 |
| | NT | 1 | 0.258 | |
| UvrC | INH | 0.185 | 0.08 | 0.01813 |
| | NT | 1 | 0.753 | |
| UvrD | INH | 0.362 | 0.136 | 0.02415 |
| | NT | 1 | 0.505 | |
| XthA | INH | 0.741 | 0.339 | 0.39897 |
| | NT | 1 | 0.282 | |
| AdnA | EMB | 1.203 | 0.141 | 0.86275 |
| | NT | 1 | 0.185 | |
| AhpC | EMB | 0.568 | 0.062 | 0.6683 |
| | NT | 1 | 0.206 | |
| AlkA | EMB | 0.844 | 0.087 | 0.72902 |
| | NT | 1 | 0.265 | |
| Dcd:dut | EMB | 1.269 | 0.128 | 0.87932 |
| | NT | 1 | 0.228 | |
| DinB1 | EMB | 0.324 | 0.073 | 0.10778 |
| | NT | 1 | 2.273 | |
| DinB2 | EMB | 1.515 | 0.173 | 0.40975 |
| | NT | 1 | 0.367 | |
| DNA ligase | EMB | 1.705 | 0.256 | 0.74016 |
| | NT | 1 | 0.159 | |

| | | | | |
|---|---|---|---|---|
| DnaE2 | EMB | 0.818 | 0.13 | 0.73481 |
| | NT | 1 | 0.497 | |
| Dut | EMB | 0.295 | 0.035 | 0.18335 |
| | NT | 1 | 0.547 | |
| End | EMB | 1.214 | 0.127 | 0.30429 |
| | NT | 1 | 0.489 | |
| KatG1 | EMB | 0.237 | 0.038 | 0.23333 |
| | NT | 1 | 0.71 | |
| LexA | EMB | 2.047 | 0.193 | 0.80759 |
| | NT | 1 | 0.193 | |
| Mfd | EMB | 1.12 | 0.141 | 0.9215 |
| | NT | 1 | 0.234 | |
| Mpg | EMB | 1.054 | 0.279 | 0.73747 |
| | NT | 1 | 0.631 | |
| MutM1 | EMB | 0.268 | 0.043 | 0.06411 |
| | NT | 1 | 0.42 | |
| MutT1 | EMB | 0.463 | 0.059 | 0.54076 |
| | NT | 1 | 0.567 | |
| MutT2 | EMB | 0.311 | 0.057 | 0.2705 |
| | NT | 1 | 1.246 | |
| MutT3 | EMB | 0.243 | 0.034 | 0.26496 |
| | NT | 1 | 0.758 | |
| MutT4 | EMB | 1.004 | 0.147 | 0.96784 |
| | NT | 1 | 0.398 | |
| MutY | EMB | 0.84 | 0.081 | 0.77104 |
| | NT | 1 | 0.356 | |
| Nei1 | EMB | 0.175 | 0.043 | 0.18413 |
| | NT | 1 | 2.692 | |
| Nei2 | EMB | 0.305 | 0.035 | 0.28952 |
| | NT | 1 | 1.048 | |
| NucS | EMB | 1.133 | 0.113 | 0.7434 |
| | NT | 1 | 0.263 | |
| Ogt | EMB | 1.84 | 0.19 | 0.42958 |
| | NT | 1 | 0.19 | |
| PolA | EMB | 1.064 | 0.116 | 0.85379 |
| | NT | 1 | 0.227 | |
| RecA | EMB | 1.413 | 0.154 | 0.08018 |
| | NT | 1 | 0.286 | |
| RecX | EMB | 1.665 | 0.281 | 0.73135 |
| | NT | 1 | 0.142 | |
| TagA | EMB | 0.902 | 0.101 | 0.83606 |
| | NT | 1 | 0.294 | |
| ThyA | EMB | 0.202 | 0.024 | 0.31757 |
| | NT | 1 | 0.688 | |
| ThyX | EMB | 0.526 | 0.071 | 0.40416 |
| | NT | 1 | 0.342 | |

| | | | | |
|---|---|---|---|---|
| UdgB | EMB | 3.51 | 0.378 | 0.33896 |
| | NT | 1 | 0.18 | |
| UdgX | EMB | 0.431 | 0.057 | 0.08724 |
| | NT | 1 | 0.599 | |
| Ung | EMB | 2.938 | 0.618 | 0.79681 |
| | NT | 1 | 0.109 | |
| UvrA | EMB | 0.878 | 0.114 | 0.82035 |
| | NT | 1 | 0.378 | |
| UvrB | EMB | 1.129 | 0.134 | 0.98668 |
| | NT | 1 | 0.316 | |
| UvrC | EMB | 0.47 | 0.061 | 0.29949 |
| | NT | 1 | 0.434 | |
| UvrD | EMB | 0.174 | 0.025 | 0.13545 |
| | NT | 1 | 0.627 | |
| XthA | EMB | 1.478 | 0.287 | 0.76146 |
| | NT | 1 | 0.245 | |
| AdnA | RIF | 2.277 | 0.204 | 0.1663 |
| | NT | 1 | 0.122 | |
| AhpC | RIF | 0.228 | 0.029 | 0.04564 |
| | NT | 1 | 0.272 | |
| AlkA | RIF | 2.122 | 0.14 | 0.63226 |
| | NT | 1 | 0.115 | |
| Dcd:dut | RIF | 0.665 | 0.083 | 0.66736 |
| | NT | 1 | 0.266 | |
| DinB1 | RIF | 0.695 | 0.177 | 0.48363 |
| | NT | 1 | 0.273 | |
| DinB2 | RIF | 1.379 | 0.138 | 0.15913 |
| | NT | 1 | 0.129 | |
| DNA ligase | RIF | 0.958 | 0.139 | 0.12799 |
| | NT | 1 | 0.109 | |
| DnaE2 | RIF | 0.66 | 0.028 | 1 |
| | NT | 1 | 0.184 | |
| Dut | RIF | 0.878 | 0.046 | 0.87121 |
| | NT | 1 | 0.109 | |
| End | RIF | 0.487 | 0.105 | 0.54775 |
| | NT | 1 | 0.527 | |
| KatG1 | RIF | 0.806 | 0.063 | 0.95255 |
| | NT | 1 | 0.113 | |
| LexA | RIF | 0.848 | 0.37 | 0.71664 |
| | NT | 1 | 0.357 | |
| Mfd | RIF | 1.38 | 0.075 | 0.6808 |
| | NT | 1 | 0.105 | |
| Mpg | RIF | 1.369 | 0.08 | 0.03145 |
| | NT | 1 | 0.192 | |
| MutM1 | RIF | 0.752 | 0.027 | 0.74627 |
| | NT | 1 | 0.124 | |

| | | | | |
|---|---|---|---|---|
| MutT1 | RIF | 0.895 | 0.088 | 0.79417 |
| | NT | 1 | 0.113 | |
| MutT2 | RIF | 0.395 | 0.035 | 0.11781 |
| | NT | 1 | 0.343 | |
| MutT3 | RIF | 0.392 | 0.064 | 0.26692 |
| | NT | 1 | 0.374 | |
| MutT4 | RIF | 2.146 | 0.337 | 0.91444 |
| | NT | 1 | 0.181 | |
| MutY | RIF | 0.894 | 0.132 | 0.66026 |
| | NT | 1 | 0.226 | |
| Nei1 | RIF | 0.744 | 0.079 | 0.68867 |
| | NT | 1 | 0.443 | |
| Nei2 | RIF | 0.751 | 0.119 | 0.65362 |
| | NT | 1 | 0.303 | |
| NucS | RIF | 0.929 | 0.046 | 0.81321 |
| | NT | 1 | 0.147 | |
| Ogt | RIF | 3.192 | 0.422 | 0.18641 |
| | NT | 1 | 0.098 | |
| PolA | RIF | 1.683 | 0.298 | 0.67136 |
| | NT | 1 | 0.118 | |
| RecA | RIF | 0.754 | 0.043 | 0.48499 |
| | NT | 1 | 0.162 | |
| RecX | RIF | 0.598 | 0.055 | 0.11886 |
| | NT | 1 | 0.158 | |
| TagA | RIF | 0.804 | 0.094 | 0.79958 |
| | NT | 1 | 0.208 | |
| ThyA | RIF | 0.462 | 0.041 | 0.5 |
| | NT | 1 | 0.172 | |
| ThyX | RIF | 0.421 | 0.061 | 0.5 |
| | NT | 1 | 0.362 | |
| UdgB | RIF | 1.687 | 0.195 | 0.25647 |
| | NT | 1 | 0.11 | |
| UdgX | RIF | 3.289 | 0.129 | 0.55937 |
| | NT | 1 | 0.088 | |
| Ung | RIF | 0.993 | 0.174 | 0.97323 |
| | NT | 1 | 0.108 | |
| UvrA | RIF | 1.39 | 0.056 | 0.83498 |
| | NT | 1 | 0.133 | |
| UvrB | RIF | 1.453 | 0.098 | 0.8279 |
| | NT | 1 | 0.103 | |
| UvrC | RIF | 0.423 | 0.027 | 0.12612 |
| | NT | 1 | 0.266 | |
| UvrD | RIF | 0.771 | 0.029 | 0.69513 |
| | NT | 1 | 0.093 | |
| XthA | RIF | 1.035 | 0.053 | 0.90709 |
| | NT | 1 | 0.103 | |

**Table S5** Oligonucleotides used for the dNTP measurements

| Name | Sequence (5'→3') |
|---|---|
| NDP-1 primer | CCGCCTCCACCGCC |
| FAM-dTTP probe | **6-FAM/**AGGACCGAG**/ZEN/**GCAAGAGCGAGCGA**/IBFQ** |
| FAM-dATP probe | **6-FAM/**TGGTCCGTG**/ZEN/**GCTTGTGCGTGCGT**/IBFQ** |
| FAM-dGTP probe | **6-FAM/**ACCATTCAC**/ZEN/**CTCACACTCACTCC**/IBFQ** |
| FAM-dCTP probe | **6-FAM/**AGGATTGAG**/ZEN/**GTAAGAGTGAGTGG**/IBFQ** |
| dTTP-DT1 template | TCGCTCGCTCTTGCCTCGGTCCTTT**A**TTTGGCGGTGGAGGCGG |
| dATP-DT1 template | ACGCACGCACAAGCCACGGACCAAA**T**AAAGGCGGTGGAGGCGG |
| dCTP-DT1 template | CCACTCACTCTTACCTCAATCCTTT**G**TTTGGCGGTGGAGGCGG |
| dGTP-DT2 template | GGAGTGAGTGTGAGGTGAATGGTTT**C**TTT**C**TTTGGCGGTGGAGGCGG |

**Table S6** dNTP concentrations in cellular extracts upon treatment with drugs

| Treatment | dNTP | Sample | μM | +/-SEM | Fold change | p-value (t-probe) |
|---|---|---|---|---|---|---|
| COMBO | dGTP | Treated | 524 | 46 | 0.54 | 0.125 |
| | | Non-treated | 977 | 237 | | |
| | dCTP | Treated | 656 | 69 | 0.98 | 0.925 |
| | | Non-treated | 671 | 155 | | |
| | dTTP | Treated | 756 | 73 | 1.10 | 0.443 |
| | | Non-treated | 658 | 123 | | |
| | dATP | Treated | 231 | 71 | 0.39 | 0.161 |
| | | Non-treated | 587 | 175 | | |
| INH | dGTP | Treated | 599 | 235 | 0.40 | 0.020 |
| | | Non-treated | 1508 | 55 | | |
| | dCTP | Treated | 311 | 25 | 0.68 | 0.246 |
| | | Non-treated | 460 | 108 | | |
| | dTTP | Treated | 458 | 130 | 0.61 | 0.194 |
| | | Non-treated | 748 | 156 | | |
| | dATP | Treated | 403 | 99 | 0.54 | 0.038 |
| | | Non-treated | 748 | 87 | | |
| EMB | dGTP | Treated | 154 | 26 | 0.05 | 0.057 |
| | | Non-treated | 3025 | 675 | | |
| | dCTP | Treated | 107 | 7 | 0.16 | 0.068 |
| | | Non-treated | 686 | 234 | | |
| | dTTP | Treated | 348 | 28 | 0.20 | 0.052 |
| | | Non-treated | 1711 | 509 | | |

| | | | | | | |
|---|---|---|---|---|---|---|
| | dATP | Treated | 163 | 32 | 0.15 | 0.006 |
| | | Non-treated | 1062 | 98 | | |
| RIF | dGTP | Treated | 253 | 39 | 0.65 | 0.056 |
| | | Non-treated | 387 | 17 | | |
| | dCTP | Treated | 202 | 33 | 3.30 | 0.013 |
| | | Non-treated | 60 | 6 | | |
| | dTTP | Treated | 238 | 42 | 1.50 | 0.182 |
| | | Non-treated | 158 | 22 | | |
| | dATP | Treated | 116 | 13 | 0.96 | 0.866 |
| | | Non-treated | 121 | 25 | | |
| MMC | dGTP | Treated | 476 | 37 | 0.43 | 0.035 |
| | | Non-treated | 1103 | 198 | | |
| | dCTP | Treated | 230 | 16 | 2.20 | 0.025 |
| | | Non-treated | 107 | 27 | | |
| | dTTP | Treated | 563 | 91 | 1.30 | 0.342 |
| | | Non-treated | 432 | 91 | | |
| | dATP | Treated | 212 | 22 | 0.60 | 0.154 |
| | | Non-treated | 355 | 67 | | |
| CIP | dGTP | Treated | 587 | 21 | 0.43 | 0.002 |
| | | Non-treated | 1380 | 114 | | |
| | dCTP | Treated | 371 | 20 | 1.10 | 0.311 |
| | | Non-treated | 336 | 47 | | |
| | dTTP | Treated | 5296 | 557 | 7.00 | 0.003 |
| | | Non-treated | 757 | 87 | | |
| | dATP | Treated | 4338 | 574 | 7.10 | 0.008 |
| | | Non-treated | 612 | 69 | | |

**Table S7** Interpreting WGS data files deposited in the European Nucleotide Archive (ENA)

| Experiment | Treatment | Sample | Sample description | ENA file name | Experiment length |
|---|---|---|---|---|---|
| MA | Ciprofloxacin | CIP1 | 5 different lineages of ciprofloxacin treated Mycobacterium smegmatis | CipA.rg.bam | 60 days |
| MA | | CIP2 | 5 different lineages of ciprofloxacin treated Mycobacterium smegmatis | CipB.rg.bam | 60 days |
| MA | | CIP3 | 5 different lineages of ciprofloxacin treated Mycobacterium smegmatis | CipC.rg.bam | 60 days |
| MA | Rifampicin | RIF1 | 6 different lineages of rifampicin treated Mycobacterium smegmatis | RIFA.rg.bam | 60 days |
| MA | | RIF2 | 5 different lineages of rifampicin treated Mycobacterium smegmatis | RIFB.rg.bam | 60 days |
| MA | | RIF3 | 5 different lineages of rifampicin treated Mycobacterium smegmatis | RIFC.rg.bam | 60 days |
| MA | Isoniazid | INH1 | 5 different lineages of isoniazid treated Mycobacterium smegmatis | INH1.rg.bam | 60 days |
| MA | | INH2 | 5 different lineages of isoniazid treated Mycobacterium smegmatis | INH2.rg.bam | 60 days |
| MA | | INH3 | 5 different lineages of isoniazid treated Mycobacterium smegmatis | INHA.rg.bam | 60 days |
| MA | Ethambutol | EMB1 | 5 different lineages of ethambutol treated Mycobacterium smegmatis | EMBA.rg.bam | 60 days |
| MA | | EMB2 | 6 different lineages of ethambutol treated Mycobacterium smegmatis | EMBB.rg.bam | 60 days |
| MA | | EMB3 | 5 different lineages of ethambutol treated Mycobacterium smegmatis | EMBC.rg.bam | 60 days |
| MA | Combination of first line drugs | COMBO1 | 5 different lineages of pyrazinamide, ethambutol, isoniazid and rifampicin treated Mycobacterium smegmatis | CombA.rg.bam | 60 days |
| MA | | COMBO2 | 6 different lineages of pyrazinamide, ethambutol, isoniazid and rifampicin treated Mycobacterium smegmatis | CombB.rg.bam | 60 days |
| MA | | COMBO3 | 5 different lineages of pyrazinamide, ethambutol, isoniazid and rifampicin treated Mycobacterium smegmatis | CombC.rg.bam | 60 days |
| MA | MitomycinC | MMC1 | 5 different lineages of mitomycinC treated Mycobacterium smegmatis | MMCA.rg.bam | 60 days |
| MA | | MMC2 | 5 different lineages of mitomycinC treated Mycobacterium smegmatis | MMCB.rg.bam | 60 days |
| MA | | MMC3 | 5 different lineages of mitomycinC treated Mycobacterium smegmatis | MMCC.rg.bam | 60 days |
| MA | UV | UV1 | 5 different lineages of UV radiation treated Mycobacterium smegmatis | UV1.rg.bam | 60 days |
| MA | | UV2 | 5 different lineages of UV radiation treated Mycobacterium smegmatis | UV2.rg.bam | 60 days |
| MA | | UV3 | 6 different lineages of UV radiation treated Mycobacterium smegmatis | UV3.rg.bam | 60 days |
| MA | Control, no treatment | MOCK1 | 5 different lineages of untreated Mycobacterium smegmatis | MockA.rg.bam | 120 days |
| MA | | MOCK2 | 5 different lineages of untreated Mycobacterium smegmatis | MockB.rg.bam | 120 days |
| MA | | MOCK3 | 5 different lineages of untreated Mycobacterium smegmatis | MockC.rg.bam | 120 days |
| MA | | MOCK4 | 5 different lineages of untreated Mycobacterium smegmatis | MockD.rg.bam | 120 days |
| MA | | MOCK5 | 5 different lineages of untreated Mycobacterium smegmatis | MockE.rg.bam | 120 days |
| MA | | MOCK6 | 5 different lineages of untreated Mycobacterium smegmatis | MockF.rg.bam | 120 days |

| | | | | | |
|---|---|---|---|---|---|
| MA | | MOCK7 | 5 different lineages of untreated Mycobacterium smegmatis | MockG.rg.bam | 120 days |
| MA | | MOCK8 | 5 different lineages of untreated Mycobacterium smegmatis | MockH.rg.bam | 120 days |
| MA | | MOCK9 | 5 different lineages of untreated Mycobacterium smegmatis | MockI.rg.bam | 60 days |
| MA | | MOCK10 | 6 different lineages of untreated Mycobacterium smegmatis | MockJ.rg.bam | 60 days |
| MA | | MOCK11 | 5 different lineages of untreated Mycobacterium smegmatis | MockK.rg.bam | 60 days |
| MA | | MOCK12 | 5 different lineages of untreated Mycobacterium smegmatis | MockL.rg.bam | 60 days |
| MA | No treatment | APAW | 1 strain, common ancestor of every treated and untreated lineages, wild-type Mycobacterium smegmatis | APAW.rg.bam | - |
| MA | No treatment | WT_MSM | 1 strain, wild-type Mycobacterium smegmatis | WT_Msm.rg.bam | - |
| Fluctuation assay | Ciprofloxacin | A03 | 5 different lineages of 0.3 μg/ml ciprofloxacin treated Mycobacterium smegmatis | A03.rg.bam | 4 days |
| Fluctuation assay | Ciprofloxacin | A05sel | 5 different lineages of 0.5 μg/ml ciprofloxacin treated Mycobacterium smegmatis | A05sel.rg.bam | 4 days |
| Fluctuation assay | Untreated | ANT | 5 different lineages of untreated Mycobacterium smegmatis | ANT.rg.bam | 4 days |
| Fluctuation assay | Untreated | At0 | 1 strain, common ancestor line of fluctuation assay treatment for samples A03, A05sel and ANT samples[1] | At0.rg.bam | 4 days |
| Fluctuation assay | Ciprofloxacin | B03 | 5 different lineages of 0.3 μg/ml ciprofloxacin treated Mycobacterium smegmatis | B03.rg.bam | 4 days |
| Fluctuation assay | Ciprofloxacin | B05sel | 5 different lineages of 0.5 μg/ml ciprofloxacin treated Mycobacterium smegmatis | B05sel.rg.bam | 4 days |
| Fluctuation assay | Untreated | BNT | 5 different lineages of untreated Mycobacterium smegmatis | BNT.rg.bam | 4 days |
| Fluctuation assay | Untreated | Bt0 | 1 strain, common ancestor line of fluctuation assay treatment for samples B03, B05sel and BNT samples[1] | Bt0.rg.bam | 4 days |
| Fluctuation assay | Ciprofloxacin | C03 | 5 different lineages of 0.3 μg/ml ciprofloxacin treated Mycobacterium smegmatis | C03.rg.bam | 4 days |
| Fluctuation assay | Ciprofloxacin | C05sel | 5 different lineages of 0.5 μg/ml ciprofloxacin treated Mycobacterium smegmatis | C05sel.rg.bam | 4 days |
| Fluctuation assay | Untreated | CNT | 5 different lineages of untreated Mycobacterium smegmatis | CNT.rg.bam | 4 days |
| Fluctuation assay | Untreated | Ct0 | 1 strain, common ancestor line of fluctuation assay treatment for samples C03, C05sel and CNT samples[1] | Ct0.rg.bam | 4 days |

1Note that At0, Bt0 and Ct0 strains also originated from APAW strain used for MA experiment.
